# Single cell transcriptomics reveals cell type specific features of developmentally regulated responses to lipopolysaccharide between birth and 5 years

**DOI:** 10.1101/2023.05.18.541356

**Authors:** James F. Read, Michael Serralha, Jesse Armitage, Muhammad Munir Iqbal, Mark N. Cruickshank, Alka Saxena, Deborah H. Strickland, Jason Waithman, Patrick G. Holt, Anthony Bosco

## Abstract

Human perinatal life is characterized by a period of extraordinary change during which newborns encounter abundant environmental stimuli and exposure to potential pathogens. To meet such challenges, the neonatal immune system is equipped with unique functional characteristics that adapt to changing conditions as development progresses across the early years of life, but the molecular characteristics of such adaptations remain poorly understood. The application of single cell genomics to birth cohorts provides an opportunity to investigate changes in gene expression programs elicited downstream of innate immune activation across early life at unprecedented resolution. In this study, we performed single cell RNA-sequencing of mononuclear cells collected from matched birth cord blood and 5-year peripheral blood samples following stimulation (18hrs) with two well-characterized innate stimuli; lipopolysaccharide (LPS) and Polyinosinic:polycytidylic acid (Poly(I:C)). We found that the transcriptional response to LPS was constrained at birth and predominantly partitioned into classical proinflammatory gene upregulation primarily by monocytes and IFN-signaling gene upregulation by lymphocytes. Moreover, these responses featured substantial cell-to-cell communication which appeared markedly strengthened between birth and 5 years. In contrast, stimulation with Poly(I:C) induced a robust IFN-signalling response across all cell types identified at birth and 5 years. Analysis of gene regulatory networks revealed IRF1 and STAT1 were key drivers of the LPS-induced IFN-signaling response in lymphocytes with a potential developmental role for IRF7 regulation. Additionally, we observed distinct activation trajectory endpoints for monocytes derived from LPS-treated cord and 5-year blood, which was not apparent among Poly(I:C)-induced monocytes. Taken together, our findings provide new insight into the gene regulatory landscape of immune cell function between birth and 5 years and point to regulatory mechanisms relevant to future investigation of infection susceptibility in early life.

## Introduction

Newborns experience remarkable environmental change as they transition from a protected, tolerogenic intrauterine environment to the outside world with an abundance of stimuli and pathogens(1). The neonatal immune system has evolved unique functional characteristics suited to the challenges of perinatal life(2). For example, neonatal myeloid cell cytokine production is skewed to promote Th_2_ and Th_17_ responses, while those that promote Th_1_ and Type I interferon (IFN) responses are attenuated (3–7). Within the lymphoid compartment, neonatal T cells arise from distinct hematopoietic stem cell populations(8), express more broadly reactive T cell receptors(9,10), display distinct epigenetic patterns(11,12), and exhibit impaired memory capacity(13,14) compared to adult counterparts(2). Importantly, cord blood-derived T cells demonstrate enhanced responses to innate immune signals(2), including those associated with activation of the Toll-like Receptor (TLR) family(15,16). Despite these functional adaptions, newborns are nonetheless highly susceptible to developing severe disease following microbial infections.

Innate immune responses are triggered by the binding of evolutionarily conserved pathogen-associate molecular patterns (PAMPs) to germline encoded pathogen recognition receptors (PRRs) of the innate immune system. The Toll-Like Receptor (TLR) family are the most well characterized PRRs, and are expressed on the cell surface (e.g., TLRs 1, 2, 4, 5, 6) or in intracellular endosomes (e.g., TLRs 3, 7, 8)(17). Cell surface TLRs bind to bacterial cell membrane/wall components and viral proteins, whereas the intracellular TLRs bind to nucleic acids(17). Downstream signalling of TLR activation is predominantly mediated by either Myeloid differentiation factor 88 (MyD88) and/or Toll-Interleukin 1 Receptor-domain-containing adapter-inducing IFN-β (TRIF)(17,18). TLR-signaling induces subsequent effector gene expression programs following engagement of specific transcription factor (TF), including Nuclear Factor kappa-B (NF-κB) and members of the IFN Regulatory Factor family (e.g. IRF3/7)(17,18), resulting in the promotion of proinflammatory (e.g. IL-1β, IL-6) and Type I IFN/antiviral (IFNα/β) response programs, respectively(17–20). TLR4 mediates immune responses following detection of lipopolysaccharide (LPS), a cell wall component of Gram-negative bacteria. Uniquely among TLRs, TLR4 ligation triggers both the MyD88-dependent proinflammatory and TRIF-dependent Type I IFN response pathways(17). TLR3 recognises viral RNA and triggers TRIF-dependent Type I IFN production(17,18).

Innate immunity has traditionally been viewed as a first line of defence capable of responding rapidly and non-specifically to pathogen encounters that lacks memory and therefore cannot mediate resistance to reinfection. This view has been challenged by the demonstration that exposure to vaccines, infections, or microbial products can induce prolonged epigenetic and functional changes in innate immune cells that provide enhanced, non-specific protection to subsequent encounters with the same pathogen or an unrelated pathogen(21). This evolving view of early life immunity provides motivation for more studies to further our understanding of how innate immune function at birth is programmed during the first years of life, a crucial period of heightened plasticity and window of susceptibility for the development of chronic diseases. The advent of single cell genomics enables a deeper understanding of the cell type specific, stimuli specific, and age-related changes that underlie the development and regulation of innate immune function in early life. Here, we deploy these powerful tools to analyse matched cord/peripheral blood mononuclear cell (C/PBMC) samples collected at birth and five years of age from two donors to provide a unique level of insight into the developmental regulation of innate immune function at birth versus early childhood at single cell resolution.

## Materials and Methods

### Study subjects

The study was designed to assess matched birth (CBMC) and 5 years (PBMC) blood samples following LPS and Polyinosinic:polycytidylic acid (Poly(I:C)) treatment, along with matched untreated controls, from two donors (one male, one female). The samples were curated from the Childhood Asthma Study (CAS) cohort, a prospective birth cohort described previously(22). Cord blood samples were collected from healthy, full-term, singleton births. Matched 5-year samples were collected from the same donor by home visit close to their 5^th^ birthday. Ethics was approved by The University of Western Australia (reference RA/4/1/7560), and fully informed parental consent was obtained for each subject.

### In vitro cell culture and innate immune stimulation

Cryopreserved CBMC/5yr PBMC samples were thawed and stimulated with 1ng/ml LPS (Enzo Biochem) or with 50μg/ml Poly(I:C) (InvivoGen), alongside an untreated controls, and culture plates were incubated at 37°C (5% CO_2_) for 18 hours (Extended Methods). LPS is a bacterial cell wall component and a quintessential TLR4 ligand(23). Poly(I:C) is a synthetic analogue of double-stranded RNA (dsRNA) and a potent activator of TLR3 and other viral nucleic acid sensing receptors(24). Each cryopreserved sample was cultured separately so that matched stimuli/control samples were processed alongside each other in a batch. Following culture, samples were re-suspended at target concentration of 2000 cells/μl. Post-culture viability is recorded in **Table S1**.

### Library preparation and sequencing

Single cells were processed on Chromium using the Chromium Next GEM Single Cell 3’ Kit v3.1 (4 reactions, PN-1000269, 10X Genomics) on Chip G (PN-1000127, 10x Genomics) according to the manufacturer’s protocol with targeted recovery of 5,000 cells per channel. Libraries were sequenced on the NovaSeq 6000 platform in a single batch.

### Alignment and initial quality control

Raw fastq.qz files were processed with the CellRanger Toolkit (Version 6.1.1, 10x Genomics) and the Human GRCh38 genome assembly (refdata-gex-GRCh38-2020-A) was used as the reference genome. The CellRanger count pipeline was run with default parameters. CellRanger web_summary outputs were assessed with no alerts (warnings or errors) recorded for any sample. Selected CellRanger outputs are recorded in **Table S1**; briefly, this project generated (on average) 5527.17 cells per sample with 63,098.75 mean reads per cell and a median of 1980.17 genes detected per cell, as estimated by CellRanger. The raw count matrix, and corresponding barcodes and features, were used for downstream QC and analysis, and these are available via the Gene Expression Omnibus (GSE232186).

### Sample pre-processing and quality control

Count matrices were imported into the R statistical environment (version 3.6.2) for quality control (QC) and analysis. Seurat (version 3.2.0)(25) was primarily used for pre-processing and data exploration. Briefly, cells/UMIs were excluded if they had low feature counts (approximately < 2000) according to dynamic thresholding of the count distribution or if their mitochondrial gene content was greater than three median absolute deviations (MADs) above the median. Features were filtered to only those expressed in at least 0.01% of cells. Doublets were detected and removed with DoubletFinder(26) and the cell cycle phase was estimated with the CellCycleScoring function (Seurat) using known cell cycle related genes. Pre-processing and quality control metrics are recorded in **Table S1**.

### Integration, Annotation, and Dimensionality reduction

Individual pre-processed samples were integrated with Seurat(25) using default parameters. Individual cells were annotated with Azimuth(27) using the human PBMC reference data set. Level 2 cell type resolution (default) was used, and cells were excluded if they had a score less than 0.5. Dimensionality reduction (UMAP) plot coordinates from the Azimuth reference were used to show the region/cell type that corresponds to our scRNA-Seq data, and a UMAP plots were also generated from the integrated data with Seurat to display the integrated cell type clustering, as well as group variables and marker gene expression intensities.

### Differential gene expression and Pathways analysis

To identify differentially expressed genes (DEGs) between LPS/Poly(I:C) and corresponding unstimulated controls for each cell type, we employed MAST(28) via the FindMarkers function from Seurat, and included cellular detection rate, mitochondrial gene proportion, and cell cycle phase. As each donor represents a different biological sex (male, female), this variable was also included as a latent variable. This approach was selected to accommodate our small sample size (two biological donors) (See Extended Methods). Genes were considered differentially expressed if they recorded a Bonferroni-corrected *p* value less than 0.01 and an average Log_2_ fold-change in expression greater than 0.25 (upregulated) or less than -0.25 (downregulated). Significantly enriched pathways associated with DEGs between cell type/stimuli groups were identified with enrichR(29) in R to query biologically relevant annotated gene sets from the Reactome(30), KEGG(31), and Gene Ontology(32) databases.

### Pseudotime trajectory inference

We applied Monocle3(33) to infer stimuli-related activation trajectories from transitional cellular states present in the data (Extended Methods). For each analysis, only the raw counts from cells relevant to that comparison (e.g., CBMC/5yr PBMC untreated and LPS-treated monocytes) were included. Regions enriched with unstimulated controls were selected as pseudotime start points so that trajectories extended into stimuli-activated regions.

### Gene Regulatory Network (GRN) analysis and in silico perturbations

We employed CellOracle(34) (version 0.12.0) to build GRNs and identify key TF drivers and their corresponding target genes for selected cell type/stimuli groupings. For this analysis, SCANPY(35) (1.9.3) was used for pre-processing and force directed graph construction and CellOracle was run with Python (3.10.6) on Ubuntu 22.04.1 (Extended Methods). The dataset was filtered to include the top 3000 most variable genes for each comparison, and the data was normalized, log transformed and scaled with default parameters (SCANPY). Separate analyses were run from raw counts for each cell type/stimuli comparison and group specific GRNs were constructed from the base Human promoter GRN provided. CellOracle was run with standard parameters and the Monocle3-defined pseudotime values for each cell were included for analysis. GRNs were used to perform *in silico* TF perturbations of IRF1, IRF7, and STAT1 to simulate the changes in cellular states after nullifying the regulatory effects of these TFs. The scale parameter was adjusted to suit each comparison (as recommended).

### Ligand-Receptor interaction analysis

We employed CellCall(36) to identify putative ligand-receptor (L-R) communication between selected cell types following LPS- and Poly(I:C)-induced activation. For each stimuli/age comparison, the genes were filtered to the top 3000 most variable genes (compared to untreated control) and single cell profiles were restricted to selected cell types of interest. Raw counts were use as input and the transcriptional communication profile was constructed with default parameters. The LR2TF function was used to extend the analysis by capturing putatively activated TFs downstream of receiver cell receptor binding of sender cell ligands for reciprocal communication between CD14^+^ monocytes and naïve CD4^+^ T cells.

## Results

### Cord and 5yr blood-derived single cell transcriptomic profiles display age-related compositional differences

We cultured matching CBMC and 5yr PBMC samples from two donors in the presence or absence of LPS or Poly(I:C) and generated single cell transcriptomic profiles (n=12) to investigate the immune responses elicited. Following pre-processing and quality control, 57,908 single cells were included for downstream analysis (mean = 4826.7 cells per sample) with an average of 7400.4 counts and 2190.3 genes detected per cell (**Table S1**).

We employed unbiased cell type annotation with Azimuth’s reference dataset-based single cell annotation mapping(27) (**Figure 1A**). Prominent populations of CD4^+^ and CD8^+^ T cells, B cells, NK cells, and monocytes were identified among the 28 cell types detected, alongside smaller populations of hematopoietic stem and progenitor cells (HSPC) and dendritic cells (DC) (**Figure 1A**). Dimensionality reduction (UMAP) of the integrated single cell profiles demonstrated clustering according to the major immune subset (**Figure 1B**). Two small T cell groups clustered separately from the larger T cell clusters, both of which comprised CD8^+^ central memory T cells and γδT cells (**Figure 1B**). The larger of these also contained mucosal-associated invariant T (MAIT) cells and was largely composed of cells isolated from 5yr PBMC samples (**Figure 1B and S1A**). Plotting the expression intensity of canonical marker genes over the integrated UMAP coordinates confirmed Azimuth annotation of the major mononuclear cell subtypes (**Figure S1B**). Negligible cell numbers were detected for several cell types (e.g., ASDC, cDC1, plasmablasts) (**Figure 1C**), and these were not considered in downstream analysis. As expected, HSPCs were more prominent among CBMC samples and memory B cells were largely restricted to 5yr PBMC samples (**Figure 1C,D**). Regulatory T cells (Treg), γδT cells, and MAIT cells also had greater cell counts at 5 years compared to birth, although they were rare or non-existent among Poly(I:C) treated samples (**Figure 1C,D**). In addition, Poly(I:C)-stimulated samples exhibited a greater ratio of CD4^+^ central memory T cells to naïve CD4^+^ T cells (**Figure 1C,D**). Analysis of the estimated cell cycle phase indicated that the majority of monocytes were in G1 phase and the majority of HSPCs were in S phase (**Figure S1A**).

**Figure 1.**
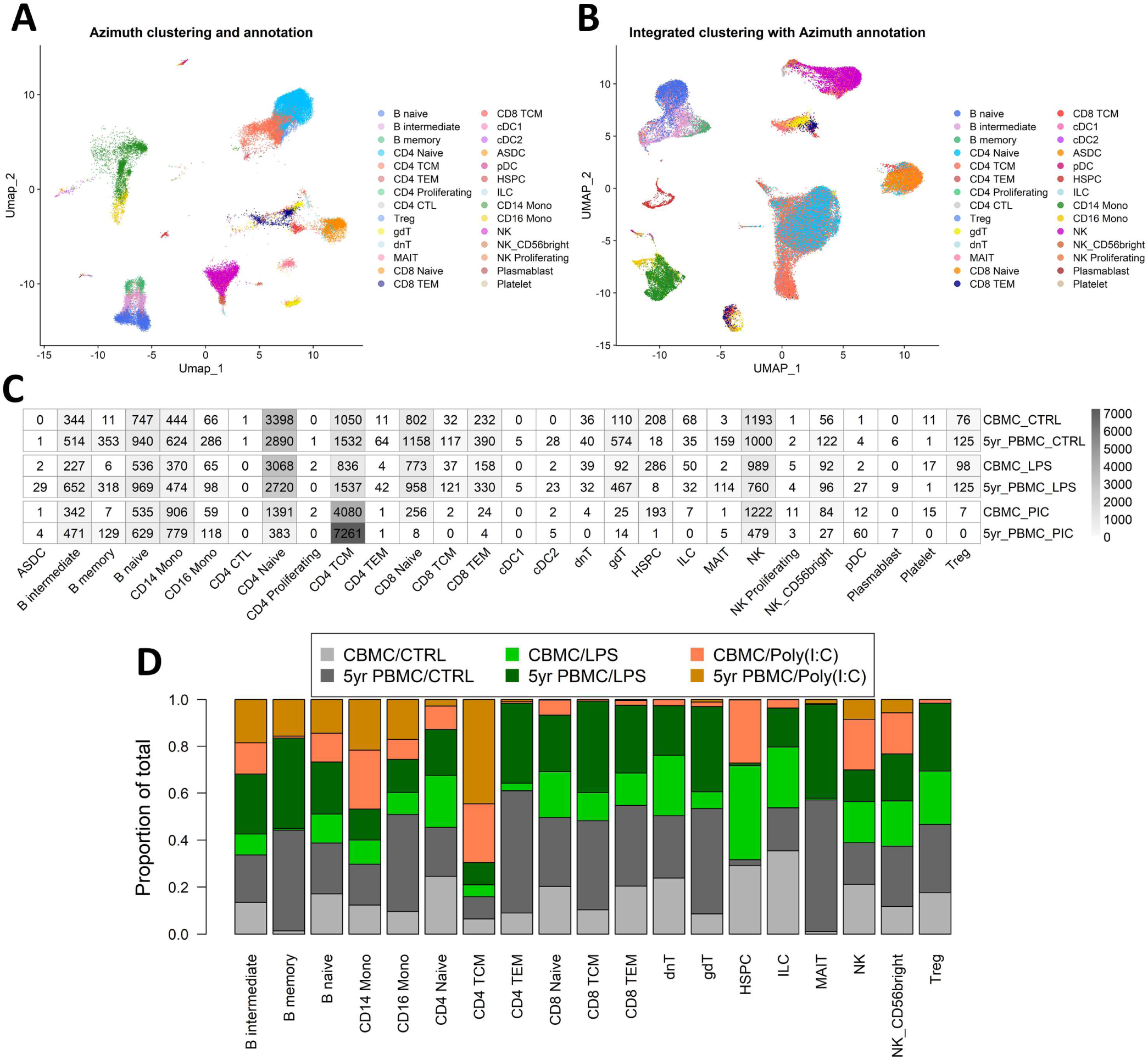
Overview of single cell RNA-Seq profiles generated. **A**) UMAP plot of scRNA-Seq data aligned to the coordinates and cell annotations of Azimuth (27). **B**) UMAP plot generated from the integrated data, overlaid with Azimuth annotations. **C**) Heatmap showing cell counts for each of the Azimuth annotated cell types stratified by the stimuli/age group. **D**) Stacked bar plot demonstrating the proportional contribution of cell numbers among stimuli/age groups.

### Differential expression analysis reveals an LPS-specific division of labour among myeloid and lymphoid immune cell compartments

To investigate changes in gene expression following *in vitro* exposure to LPS and Poly(I:C) we employed MAST(28) analysis to identify DEGs (Log_2_FC > |0.25| and Bonferroni-corrected *p* < 0.01) between stimulated cells and corresponding unstimulated controls for each cell type. In general, changes in gene expression magnitude induced by Poly(I:C) were greater than LPS for both CBMC and 5yr PBMC samples (**Figure 2A,B & S2A**). Genes encoding proinflammatory cytokines, such as *IL1B* and *CXCL8*, were prominently upregulated by monocytes following LPS stimulation of CBMCs, and this proinflammatory gene expression signature strengthen at 5 years of age (**Figure 2A-C & S2A**). Additionally, there was a marked increase in the number of DEGs by CD16^+^ monocytes between CBMC and 5yr PBMC samples (**Figure 2A,B & S2A**). Interestingly, HSPCs demonstrated a substantial gene expression response to stimuli at birth with a distinctive LPS-induced transcriptional profile – which includes *CXCL8* and *CXCL13* – compared to the IFN-related Poly(I:C)-stimulated HSPC profile (**Figure 2A & S2A**).

**Figure 2.**
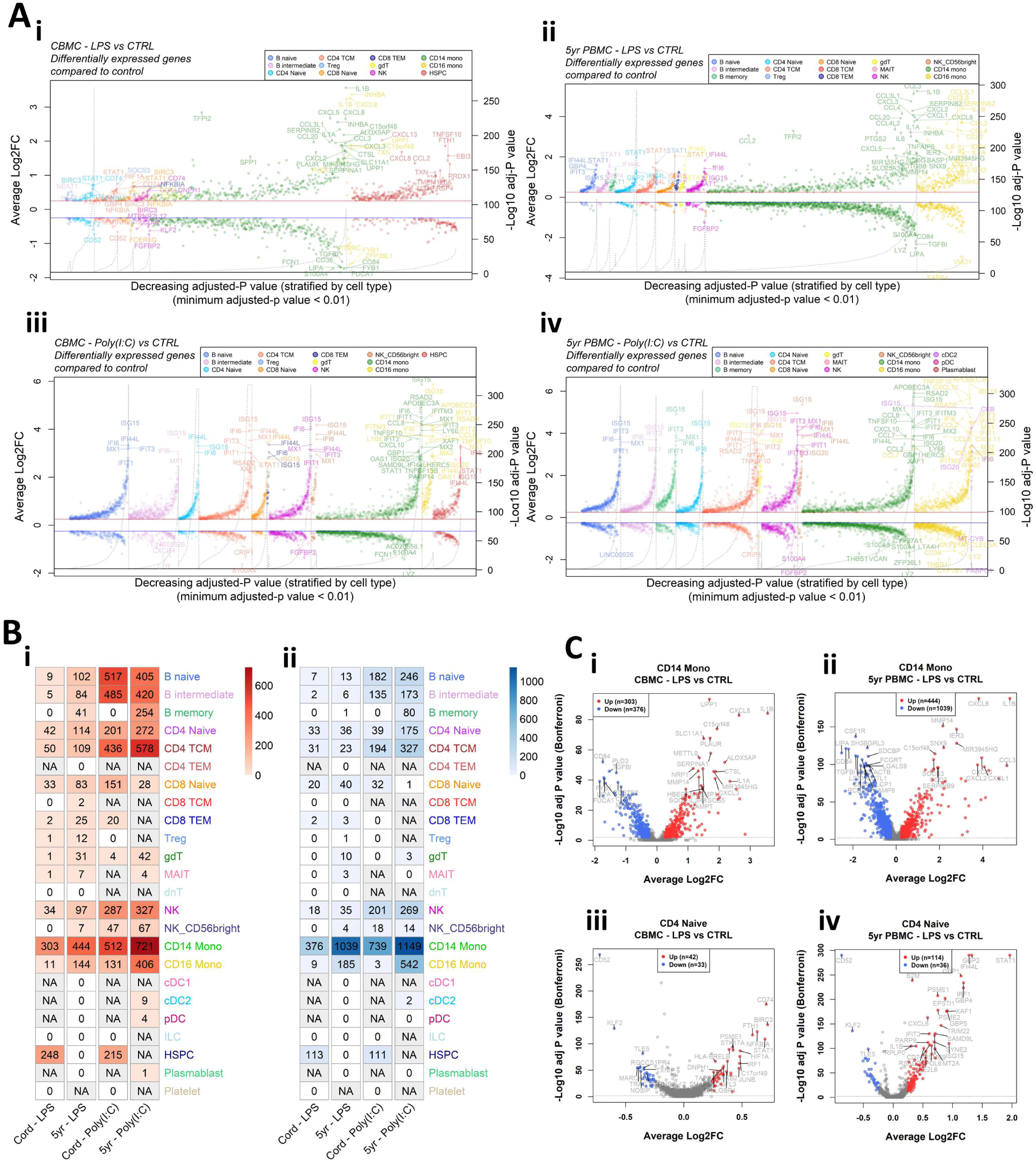
Differential expression analysis between stimulated samples and matched unstimulated controls. **A**) Aligned volcano plots displaying differentially expressed genes compared to unstimulated controls for the LPS-stimulated CBMC (**i**), LPS-stimulated 5-year PBMC (**ii**), Poly(I:C)-stimulated CBMC (**iii**), and Poly(I:C)-stimulated 5-year PBMC (**iv**) comparisons. Each point represents a differentially expressed gene (Adjusted-*p* value < 0.01 (Bonferroni correction), average Log_2_FC > 0.25) ordered by decreasing Bonferroni corrected *p* value and stratified by cell type (x-axis). The left y-axis corresponds to points on the plots and shows the average Log_2_ fold change for that gene/cell type. The right y-axis corresponds to dashed black line and represents the -Log_10_ Bonferroni corrected *p* value. Solid red line (blue line) represents an average Log_2_ fold change of 0.25 (-0.25). **B**) Heatmap showing the number of significant differentially expressed genes that were upregulated (**i**) and downregulated (**ii**) for each group compared to their corresponding unstimulated control. Columns correspond to the age/stimuli group, where cord refers to CBMC samples and 5yr refers to 5yr PBMC samples. Increased color intensity corresponds to a greater number of differentially expressed genes for that comparison and comparison with insufficient cell numbers for analysis a denoted as not applicable (grey). **C**) Volcano plots of the comparison of LPS versus unstimulated control of CD14^+^ monocytes from CBMC (**i**) and 5yr PBMC (**ii**) samples, and naïve CD4^+^ T cells from CBMC (**iii**) and 5yr PBMC (**iv**) samples. The x-axis shows the average log_2_ fold change and the y-axis shows the -Log_10_ Bonferroni-corrected *p* value. The dashed grey line indicates a Bonferroni-corrected *p* value of 0.01. Points colored red and blue represent gene with are considered significantly upregulated and downregulated, respectively.

Strikingly, several CBMC-derived T and B cell subsets exhibited upregulation of IFN-related genes, such as *IRF1* and *STAT1*, following LPS stimulation at birth (**Figure 2A,C & S2A**). Furthermore, we found that IFN-related genes exhibited enhanced upregulation in lymphocytes isolated from LPS-treated 5yr PBMCs compared to LPS-treated CBMCs (**Figure 2A-C & S2A**). Individual volcano plots of the LPS-induced gene expression comparisons for CD14^+^ monocytes and naïve CD4^+^ T cells are shown in **Figure 2C** to highlight the substantial proinflammatory monocyte response and elevated upregulation of IFN-related genes by CD4^+^ T cells at 5yrs compared to birth (**Figure 2C**). In contrast to the LPS-stimulated samples, Poly(I:C) induced a strong IFN-mediated response in all cell types in the CBMC and 5yr PBMC samples, exemplified by upregulation of IFN Stimulated Gene 15 (*ISG15*) (**Figure 2A-B, Figure S2A**). Assessment of the overlap of upregulated genes showed approximately twice as many genes were conserved between CBMC and 5yr PBMC responses to Poly(I:C) compared to LPS (**Figure S2B**). Differential gene expression results for all cell types and comparisons analysed have been compiled together and are presented in the **Supplementary data**.

The above results demonstrate that LPS treatment of mononuclear cells induces contemporaneous upregulation of classical proinflammatory genes by monocytes and IFN-related genes by lymphocytes. Furthermore, these partitioned responses show apparent strengthening between samples collected at birth and 5 years of age. Importantly, matched samples treated with Poly(I:C) demonstrated uniform upregulation of IFN-related genes across all cell types with a comparable number and change in expression magnitudes of dysregulated genes between birth and 5 years.

### IFN-signaling pathways are activated by LPS in lymphocytes at birth and intensify by 5 years

We leveraged the Reactome(30), KEGG(31), and Gene Ontology(32) databases to determine significantly enriched biological pathways from our DEG lists. LPS-induced upregulated genes from CBMC-derived naïve CD4^+^ T cells exhibited enrichment of cytokine signaling pathways, including IFN-signaling, among several immune-related pathways (**Figure 3A**). At 5 years, IFN-related pathways were among the dominant pathways identified from upregulated genes of LPS-induced naïve CD4^+^ T cells (**Figure 3A**), aligning with the findings from our differential expression analysis. Comparable results were observed for naïve B cells, naïve CD8 T cells, and NK cells (**Figure S3B-D**). Upregulated genes from LPS-induced CD14^+^ monocytes exhibited similar pathways enrichment from CBMC and 5yr PBMC samples, including several pathways associated with antibacterial responses (e.g., NF-kB and neutrophil degranulation) and specific LPS response pathways (**Figure S3A**). IFN-related pathways dominated the significantly enriched results from the upregulated gene lists for all cell types detected from Poly(I:C)-treated CBMC and 5yr PBMC samples (**Figure S4**). The pathways enrichment analyses for all cell types and comparisons (upregulated and downregulated genes) are presented in **Supplementary data**.

**Figure 3.**
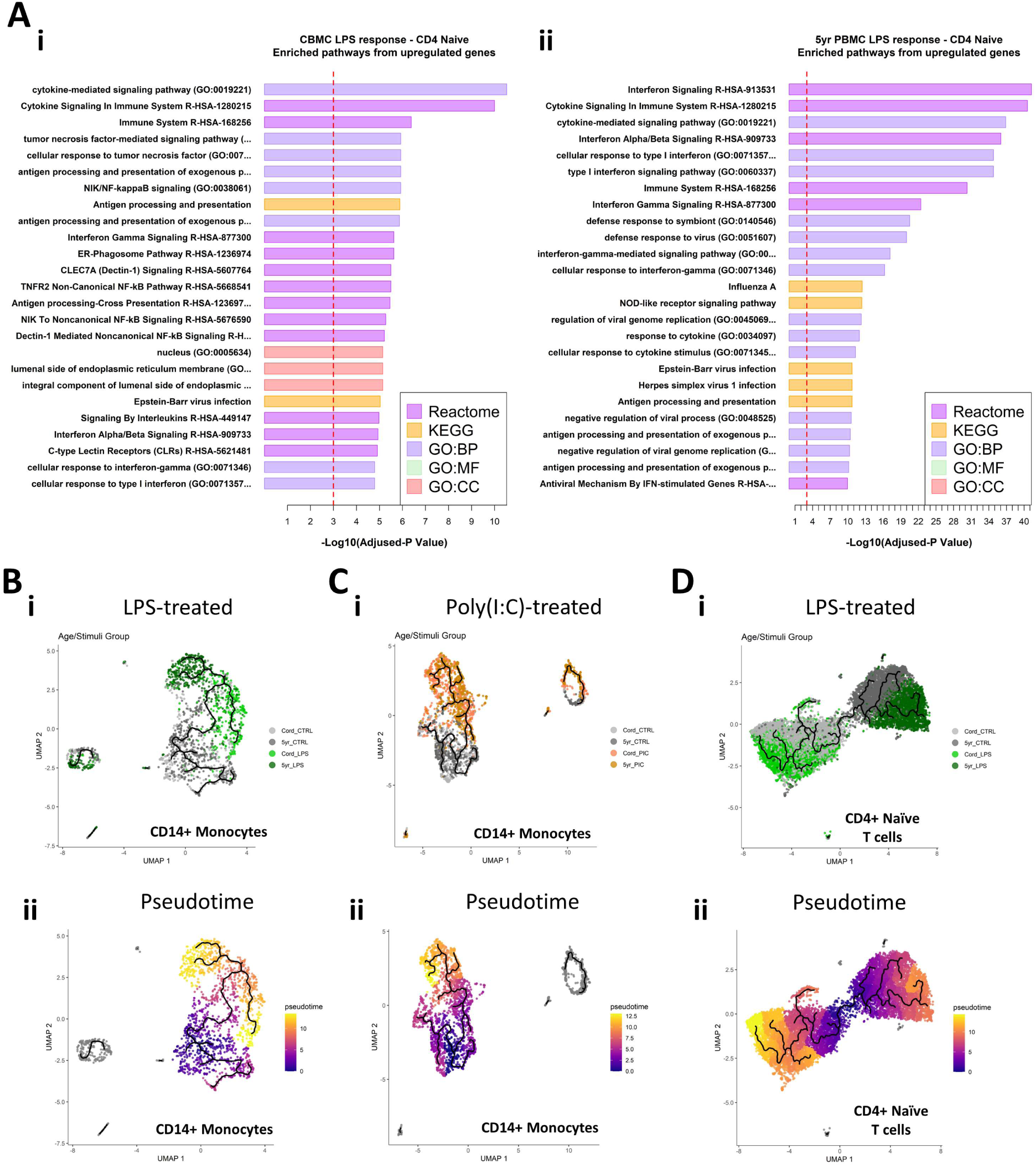
Pathway analysis of differentially expressed genes and pseudotime trajectory mapping of cellular activation trajectories. **A**) Horizontal bar plots of significantly enriched pathways from upregulated genes found for the comparison of CD4^+^ naïve T cells stimulated with LPS versus corresponding unstimulated control from CBMC (**i**) and 5yr PBMC (**ii**) samples. The x-axis shows the -Log_10_ adjusted-*p* value associated with pathways enrichment, the dashed red line indicates an adjusted-*p* value of 0.001. Results are ordered from top by decreasing adjusted-*p* value for significantly enriched pathways identified from the Reactome, KEGG, and Gene Ontology (GO) databases. BP, Biological Process; MF, Molecular Function; CC, Cellular Compartment. **B-E**) UMAP plots representing cellular activation trajectories for LPS-stimulated CD14^+^ monocytes (**B**), Poly(I:C)-stimulated CD14^+^ monocytes (**C**), and LPS-stimulated CD4^+^ naïve T cells (**D**). The first plot (**i**) in each panel is stratified by sample group and the second plot (**ii**) show the pseudotime. The branching black line on each plot represents the activation trajectory fitted to the data (monocle3 (33)).

Taken together, analysis of DEGs and pathways enrichment suggests that the immune response to Poly(I:C), and thus some viral pathogens, is to some degree hardcoded at birth to elicit a substantial IFN-mediated response from all immune cell types, and this biological feature remains relatively unchanged at 5 years of age. In contrast, LPS-induced CBMC/PBMCs demonstrate a division of labour whereby cells of the myeloid immune compartment express proinflammatory genes commonly associated with antibacterial responses at birth and age 5 whilst lymphocytes upregulate detectable quantities of IFN-signaling related genes/pathways at birth, which demonstrate a subsequent and substantial increase in prominence, and presumably function, at 5 years of age.

### Pseudotime mapping of cell differentiation trajectories demonstrates cell type- and age-specific immune activation

We next focused our analysis primarily on monocytes and naïve CD4^+^ T cells as exemplars of the myeloid and lymphoid immune compartment, respectively.

The various intermediary cellular states captured by single cell transcriptomics can be used to infer pseudotime trajectories and reconstruct biological processes(37). For this purpose, we employed monocle3(33) to infer dynamic trajectories underlying LPS and Poly(I:C) activation of CD14^+^ monocytes and naïve CD4^+^ T cells in CBMC and 5yr PBMC samples. Our analysis shows that CD14^+^ monocytes from unstimulated CBMC and 5yr PBMC control samples cluster together, indicating that baseline expression is relatively similar at birth and in early childhood (**Figure 3B**). However, stimulation with LPS promotes distinct activation endpoints with respect to the age at which the sample was collected (**Figure 3B**). In contrast, treatment with Poly(I:C) produces a single activation trajectory for CD14^+^ Monocytes isolated from CBMC and 5yr PBMC samples (**Figure 3C**). The unstimulated controls from CBMC- and 5yr PBMC-derived naïve CD4^+^ T cells cluster separately suggesting distinct baseline profiles, and subsequently produced distinct activation states following LPS treatment (**Figure 3D**). Similar trajectory characteristics were observed for CD8^+^ T, B, and NK cells stimulated with LPS (**Figure S5A**), although there was minimal distinction between LPS-treated and control B and NK cells from CBMC samples – likely explained by their relatively small gene expression change following LPS-activation. Gene expression profiles from T, B, and NK cells treated with Poly(I:C) exhibited a stimuli effect substantially greater than the difference between the CBMC and 5yr PBMC responses, resulting in the inability to fit a biologically meaningful trajectory to the data (**Figure S5B**).

### Gene regulatory network analysis identifies IRF1 and STAT1 as key regulators of LPS-induced lymphocyte response

Genes are expressed by the coordinated action of cis-regulatory elements and TFs. Construction of context specific Gene Regulatory Networks (GRN) allows investigation of the relationship between TFs and their target genes (TGs). To this end, we constructed context-specific GRNs with CellOracle(34) to identify TFs which act as the master regulators of the innate immune responses investigated in this study. This approach identified the key IFN-signaling drivers STAT1, IRF1, and IRF7 among the top TFs identified from LPS-induced naïve CD4^+^ T cell isolated from CBMC samples (**Figure 4A**). Furthermore, STAT1 and IRF1 were the top regulators of the 5yr naïve CD4^+^ T cell response to LPS, using eigenvector centrality as the metric. Indeed, these TFs were among the top regulators of all LPS-induced lymphocytes assessed in this study from CBMC and 5yr PBMC samples, and this result was consistent across metrics used to assess the GRN (eigenvector, betweenness, or degree centrality) (**Figure S6A-D**). Several known mediators of myeloid inflammation (e.g., FOS, JUN, ATF3) were identified as the key regulators of the LPS-induced monocyte responses (**Figure S6E**).

**Figure 4.**
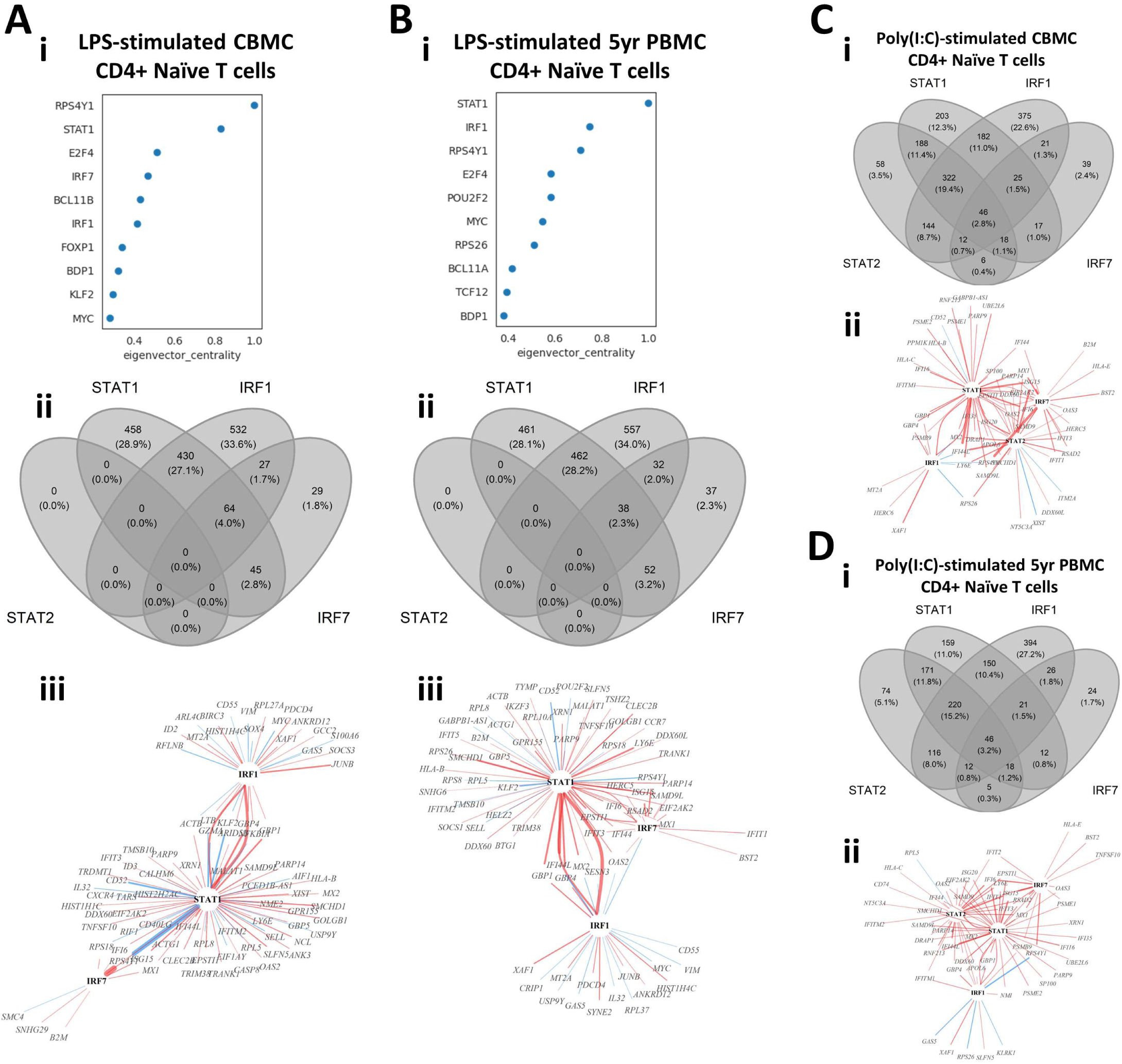
Identification of master regulators of *in vitro* stimulation responses of CBMC- and 5yr PBMC-derived naïve CD4^+^ T cells. **A-B**) Output plots from CellOracle (34) network analysis showing transcription factors and target genes ranked by eigenvector centrality for LPS-induced naïve CD4^+^ T cells (**i**, **ii**) isolated from CBMC (**A**) and 5yr PBMC (**B**) samples (panels continue downward). Venn diagrams (**ii**) depict the overlap in target genes between the selected IFN-related transcription factors identified in the CellOracle analysis for the corresponding analysis above; all target genes which were significantly associated (*p* value < 0.01) with one of the IFN-related TFs were included in the analysis. TF-TG wiring diagrams (**iii**) illustrate the interrelationship between the selected IFN-related TF and the top 100 connections to target genes by connection strength (absolute coefficient) for the corresponding analysis above. The regulators (TFs) are identified by black text and target genes are identified by grey text. Red connections indicate positive relationships (increased TG activity) and blue connections indicate a negative relationships (suppressed TG activity). The width of the connecting line illustrates relative strength with the network shown. **C-D**) Venn diagrams (**i**) and TF-TG wiring diagrams (**ii**) for the analysis of Poly(I:C)-induced naïve CD4^+^ T cells isolated from CBMC (**C**) and 5yr PBMC (**D**) samples. Analysis and plotting parameters are identical to those used for the LPS-induced response shown in A and B.

As we previously identified IFN-related genes and pathways characterized the lymphocyte response to LPS, we further investigated the role of IRF1, IRF7, and STAT1, alongside related TF STAT2, within our experimental context. We used Venn diagrams to visualize the overlap of all target genes with significant connections (*p* value < 0.01) to these TFs (**Figure 4A,B**). This analysis demonstrated similar overlap in target genes between CBMC- and 5yr PBMC-derived naïve CD4^+^ T cells stimulated with LPS. Additionally, IRF1 and STAT1 accounted for approximately 90% of TG connections and the response network was independent of STAT2 activity (**Figure 4A,B**). We next plotted wiring diagrams of the top 100 TG connections for the selected TFs to assess the relationships among the most strongly regulated interactions (**Figure 4A,B**). IRF1 and STAT1 were strongly connected to each other, reflecting the functional association of IRF1 with STAT1 homodimers(38), and STAT1 demonstrated the greatest number of connections for LPS-induced naïve CD4^+^ T cells birth and 5 years (**Figure 4A,B**). Interestingly, IRF7, which was only peripherally connected at birth, integrates more strongly with STAT1 and IRF1 via IFN-related TGs (e,g., *ISG15*, *IFIT3*, *IFI6*) at 5 years (**Figure 4A,B**). IRF1, IRF7, STAT1, and STAT2 were consistently among the top drivers of the Poly(I:C)-induced lymphocyte response following identical analysis (**Figure S7A-D**). The overlap of significantly connected TG was similar between Poly(I:C)-induced naïve CD4^+^ T cells from CBMC and 5yr PBMC samples and the contribution of STAT2 stands in stark contrast to its absence in the corresponding LPS response analysis (**Figure 4C,D**). Additionally, wiring diagrams of the most connected TG were dominated by the interaction between IRF7, STAT1, and STAT2 and IFN-associated target genes, and this was comparable between birth and 5 years of age (**Figure 4C,D**).

A core functionality of CellOracle is the ability to perform *in silico* perturbations to simulate TF knock-out and assess the outcome on cellular states(34). Simulated knock-out of *IRF1* or *STAT1* results in a reversal of the activation trajectory of LPS-induced naïve CD4^+^ T cells from CBMC and, to a lesser extent, 5yr PBMC samples, whereas knock-out of IRF7 primarily affects 5yr PBMC-derived samples (**Figure S8**), aligning with inferences from the corresponding wiring diagrams (**Figure 4A,B**). These findings demonstrate that IRF1 and STAT1 are central drivers of the lymphocyte response to LPS in early life, and suggest an important developmental role for IRF7.

### Ligand-receptor interaction analysis reveal dynamic immune cell crosstalk patterns at birth and 5 years

PAMP recognition by PRRs provokes downstream production of effector ligands by activated immune cells. These ligands enact subsequent intercellular communication by binding to their cognate receptors on other immune cells, prompting a signal cascade which mediates downstream transcription factor activity. We employed CellCall(36) to identify putative cell-cell communication pairings and capture signaling cascades following LPS- or Poly(I:C)-induced activation of selected cell types in this study. This approach revealed complex interconnectedness within lymphoid subsets and between lymphoid subsets and monocytes from stimulated samples (**Figure 5A**). HSPCs also demonstrated ligand-receptor pairing (**Figure 5A**), suggesting they participate in the CBMC response to stimuli investigated in this study. Interestingly, receptors of several 5yr PBMC-derived lymphocyte subsets (e.g., naïve CD4^+^ and CD8^+^ T cells) recorded limited or absent incoming signals, which were active in corresponding CBMC-derived samples (**Figure 5A**).

**Figure 5.**
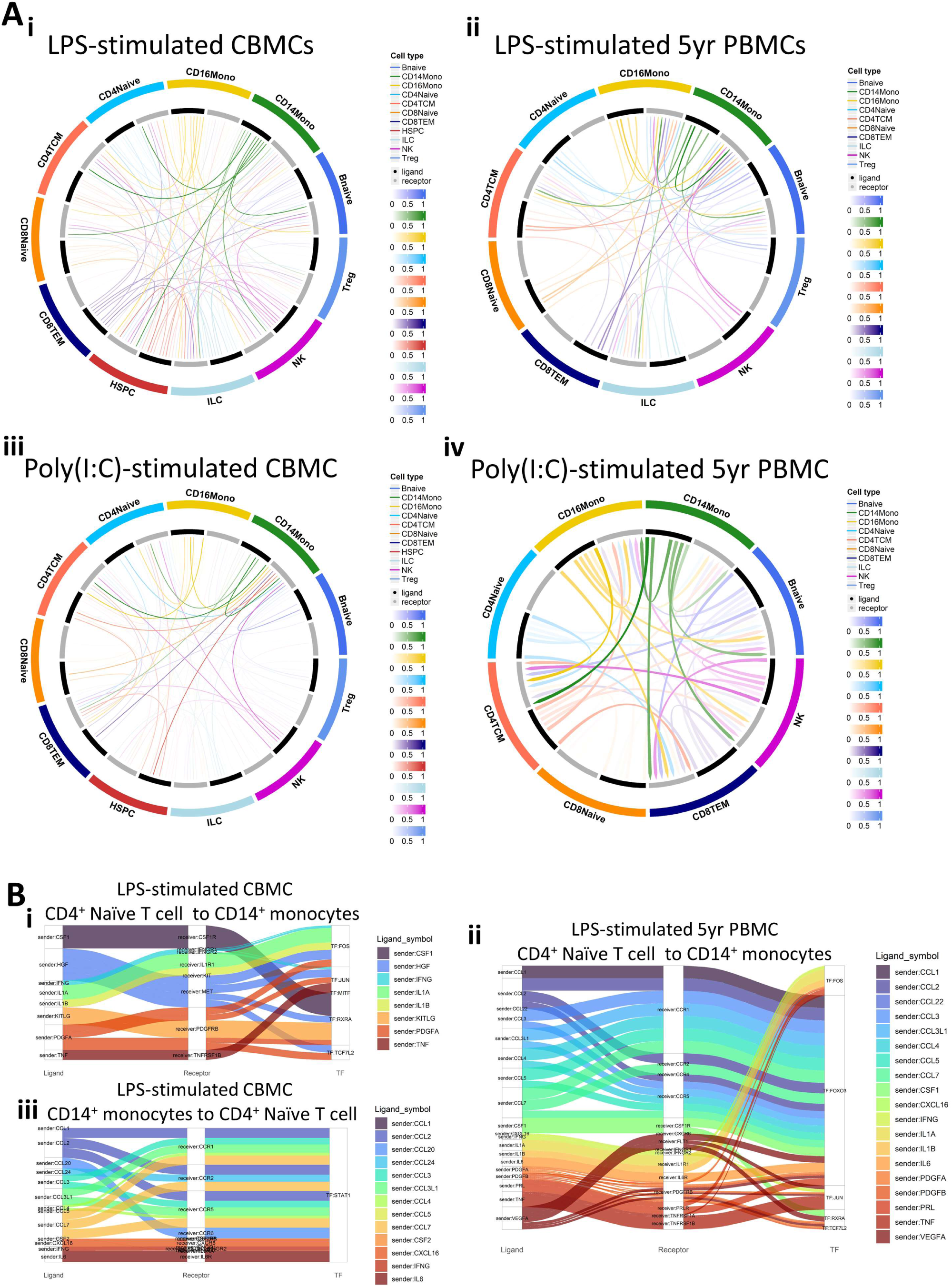
Cell-to-cell communication and subsequent transcription factor activation. **A**) Circos plots of ligand-receptor communication for selected cell types for LPS-stimulated CBMC (**i**) and 5yr PBMC (**ii**) samples and Poly(I:C)-stimulated CBMC (**iii**) and 5yr PBMC (**iv**) samples. Cell type ligands are represented by black bars, cell type receptors are represented by grey bars, and the strength of the signal is represented by the color intensity. **B**) Sankey plots demonstrating the cell-to-cell communication from ‘sender’ cell ligands (left column) to ‘receiver’ cell receptors (central column) and the subsequent transcription factor putatively activated from the ligand-receptor interaction (right column). Plot shows the communication from naïve CD4^+^ T cells to CD14^+^ monocytes for LPS-induced CBMC (**i**) and 5yr PBMC (**ii**) samples, and the reverse communication (CD14^+^ monocytes to naïve CD4^+^ T cells) for LPS-induced CBMC samples (**iii**). TF; Transcription factor.

We next focussed on the receptor-ligand interactions between LPS-induced naïve CD4^+^ T cells and CD14^+^ monocytes. At birth, CD4^+^ T cell-associated cytokine and growth factor ligands (e.g., IFNG, IL1A/B, PDGFA) were identified as inducers of several TFs, including the activating protein-1 (AP-1) TF complex members FOS and JUN (**Figure 5B(i)**). JUN and FOS are also predicted to be activated by naïve CD4^+^ T cell ligands in CD14^+^ monocytes at 5 years, and there is prominent additional signaling from suite of chemokine (e.g., CCL1-5) promoting FOXO3 activity (**Figure 5B(ii)**). Analysis of the reverse direction identified interactions between CD14^+^ monocytes ligands, including CCL chemokines, IFNG, and IL6, and naïve CD4^+^ T cell receptors, all of which putatively induced STAT1 activity which may in part explain the STAT1 prominence within the LPS-induced naïve CD4^+^ T cell regulatory networks observed in **Figure 4**. No interactions from CD14^+^ monocyte ligands to naïve CD4^+^ T cell receptors were detected for 5-year PBMC samples from this analysis. Taken together, the intracellular signaling analyses reveal complex ligand-receptor interactions between immune cell types isolated from CBMC and 5yr PBMC samples treated with LPS and Poly(I:C), and that the relationship of these interactions are subject to change between birth and 5 years of age.

## Discussion

The neonatal immune system exhibits unique functional characteristics that are tailored to the challenges of perinatal life(39). Here, we employed single cell RNA-Seq to deeply profile innate immune responses to Poly(I:C) and LPS in longitudinally matched CBMC/PBMC samples collected from two donors at birth versus age five years. We found that Poly(I:C) induced a robust response across all cell types regardless of age. In contrast, LPS responses were heavily constrained at birth at which point they were largely restricted to monocytes and HSPCs. Moreover, we observed a division of labour in the LPS responses, where monocytes/HSPC upregulated proinflammatory molecules whereas lymphocyte populations elicited IRF1/STAT1-mediated IFN-signaling pathways. Importantly, these responses exhibited substantial intracellular crosstalk and markedly strengthened between birth and age 5 years. Finally, we observed distinct activation/response trajectory endpoints for CBMC- and PBMC-derived monocytes stimulated with LPS, and this was not apparent among samples exposed to Poly(I:C). Despite the size of our study, these findings offer proof-of-principle feasibility to capture cell type-specific and context-specific gene regulatory programs that underlie innate immune function at birth versus age 5 and provides a framework for future studies to track innate immune function across early life in relation to environment exposures and disease risk.

From our gene expression analysis, we found that Poly(I:C) provoked a robust IFN-signalling response by all cell types detected, and this was relatively stable between birth and 5 years. In contrast, the LPS response was more constrained at birth compared to early childhood, and proinflammatory responses were primarily mediated by CD14^+^ monocytes, alongside HSPCs, supporting their role as immune effectors(40). Strikingly, we observed a partitioning among immune cell types following LPS stimulation whereby archetypal proinflammatory genes (e.g., *IL1B*, *CXCL8*) were upregulated in CD14^+^ monocytes while IFN-signaling genes (e.g., *STAT1*) were upregulated in lymphocytes (T and B cells). Notably, the transcriptional response to LPS displayed more profound differences between birth and 5 years compared to matched samples exposed to Poly(I:C). At the level of gene regulatory networks, we found that IFN-related IRF1 and STAT1 transcription factors(41) were the master regulators of lymphocyte response to LPS stimulation, in agreement with their corresponding gene expression, whereas the LPS-induced monocyte response was mediated by inflammatory regulators (e.g., FOS, JUN, ATF3)(42,43). Furthermore, *in silico* perturbation(34) to simulate the blockade of IRF1, IRF7, and STAT1, reinforced the central control of IRF1 and STAT1 in LPS-induced IFN-signalling networks of CD4 T cells at birth, and demonstrated that the influence of IRF7, the quintessential driver of type I IFN responses(44), within this network is more pronounced at 5 years.

Immune responses are mediated by the activation of multiple cell populations that transition through dynamic molecular states. Employing pseudotime trajectory inference, we found that naïve CD4^+^ T cells from resting CBMC samples clustered separately, consistent with notion that neonatal T cells represent a distinct lineage of cells(2,8). In contrast, LPS-induced monocyte activation trajectories started from a common baseline which subsequently diverged into age-specific end points. Monocytes trajectories from Poly(I:C)-induced CBMC/5yr PBMC responses exhibited a single trajectory, suggesting that the developmental regulation of innate immune function is much more profound for LPS responses compared with Poly(I:C) responses, as we have reported previously(45,46). We also observed striking differences in cellular composition between CBMC/PBMC samples, such as the lack of MAIT cells at birth, which is presumably explained by the fact that MAIT cells are driven by bacterial metabolites(47,48), and consequently develop in parallel with the infant microbiome(49).

Immune responses are governed by complex cellular interactions that are primarily mediated by ligand-receptor signals(50). We systematically inferred ligand-receptor mediated cell communication networks from the data which revealed substantial crosstalk between myeloid and lymphoid lineage cell types, indicating a coordinated response to LPS and Poly(I:C) stimulation. Additionally, we identified several age-specific interactions, such as extensive HSPC crosstalk in CBMC samples and less discriminate ligand binding to T cell surface receptors at birth. For example, molecular interactions were observed between LPS-induced monocyte ligands (e.g., IL6, CCL2/3/7) and naïve CD4^+^ T cell receptors (e.g., IL6R, CCR1/2/5) in CBMC samples that were not observed at age 5yrs. These temporally restricted monocyte-CD4^+^ T cell interactions in turn elicited STAT1 activity, suggesting that the mechanisms that determine IFN responses to LPS are qualitatively different at birth versus 5 years of age. Conversely, our analysis of naïve CD4^+^ T cell ligand interactions with monocyte receptors showed that regulatory activity for FOXO3 – which has been implicated in inflammation cytokine production and TLR4 upregulation specifically in LPS-induced monocytes(51) – was restricted to the 5-year LPS response, serving as an example of age-related differences in the regulation of LPS-induced monocyte responses that result in activation trajectory branching.

Our study has several limitations which we acknowledged, as follows. First, findings from our study are based on data collected from 12 scRNA-Seq samples generated from two biological donors, and accordingly follow-up studies in larger sample sizes are required to extend the findings to the general population. Second, we studied innate immune function at a single time point using two ligands. Future studies could investigate multiple time points and a broader panel of stimuli including intact viruses and bacteria. Additionally, we acknowledge that the immune system undergoes dynamic changes in the first week of life(52), and accordingly, innate immune responses in cord blood represent baseline responses at birth. Finally, future studies employing whole blood samples could extend these finding to cell types not captured among the mononuclear cell compartment from the samples investigated in this study. Notwithstanding these limitations, our study highlights several cell type-, stimulus-, and age-specific features of innate function of immune cells in early life, including underlying gene expression response programs, gene regulatory networks, and patterns of intercellular communication. These findings are relevant to future studies designed to dissect these mechanisms in relation to environmental exposures and disease risk.

## Data availability statement

The dataset presented in this study is available from the NCBI Gene Expression Omnibus repository with accession number GSE232186. The genes and pathways list (and corresponding statistical metrics) identified from differential expression and pathways analysis, respectively, for each cell type assessed are available in the **Supplementary Data** file.

## Ethics statement

Ethics was approved by The University of Western Australia (reference RA/4/1/7560). Written informed consent to participate in this study was provided by the participants’ legal guardian/next of kin.

## Author contributions

PGH and AB conceived the study and designed the experiments. JFR and MS assisted with the experimental design and optimized and performed the experiments. JFR, AB, PGH, MS, JA, DHS, MC, and JW were involved in determining the analysis approach and optimization. MMI and AS generated scRNA-Seq libraries and conducted read alignment. JFR analysed the data. JFR, AB, and PGH interpreted the data with assistance from JA, DHS, MC, and JW. JFR and AB drafted the manuscript. All authors contributed to the article and approved the submitted version.

## Acknowledgements

The authors would like to thank the study participants and their families for their involvement. Library preparation and Sequencing was conducted in the Genomics WA Laboratory in Perth, Australia. This facility is supported by BioPlatforms Australia, State Government Western Australia, Australian Cancer Research Foundation, Cancer Research Trust, Harry Perkins Institute of Medical Research, Telethon Kids Institute and the University of Western Australia. We gratefully acknowledge the Australian Cancer Research Foundation and the Centre for Advanced Cancer Genomics for making available Illumina Sequencers for the use of Genomics WA.

## Supplementary Table and Figures

**Table S1.**
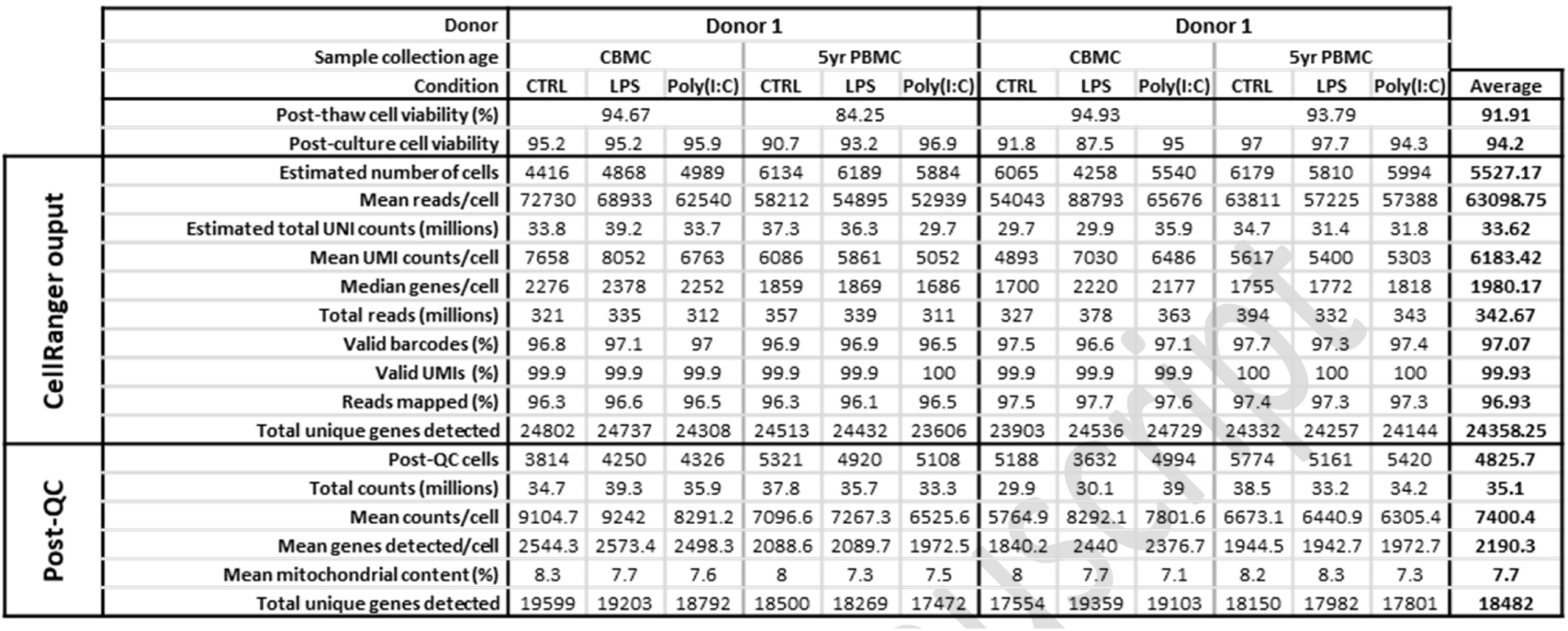
Table of quality control metrics recorded for cell viability, CellRanger, and post-alignment quality control.

**Figure S1.**
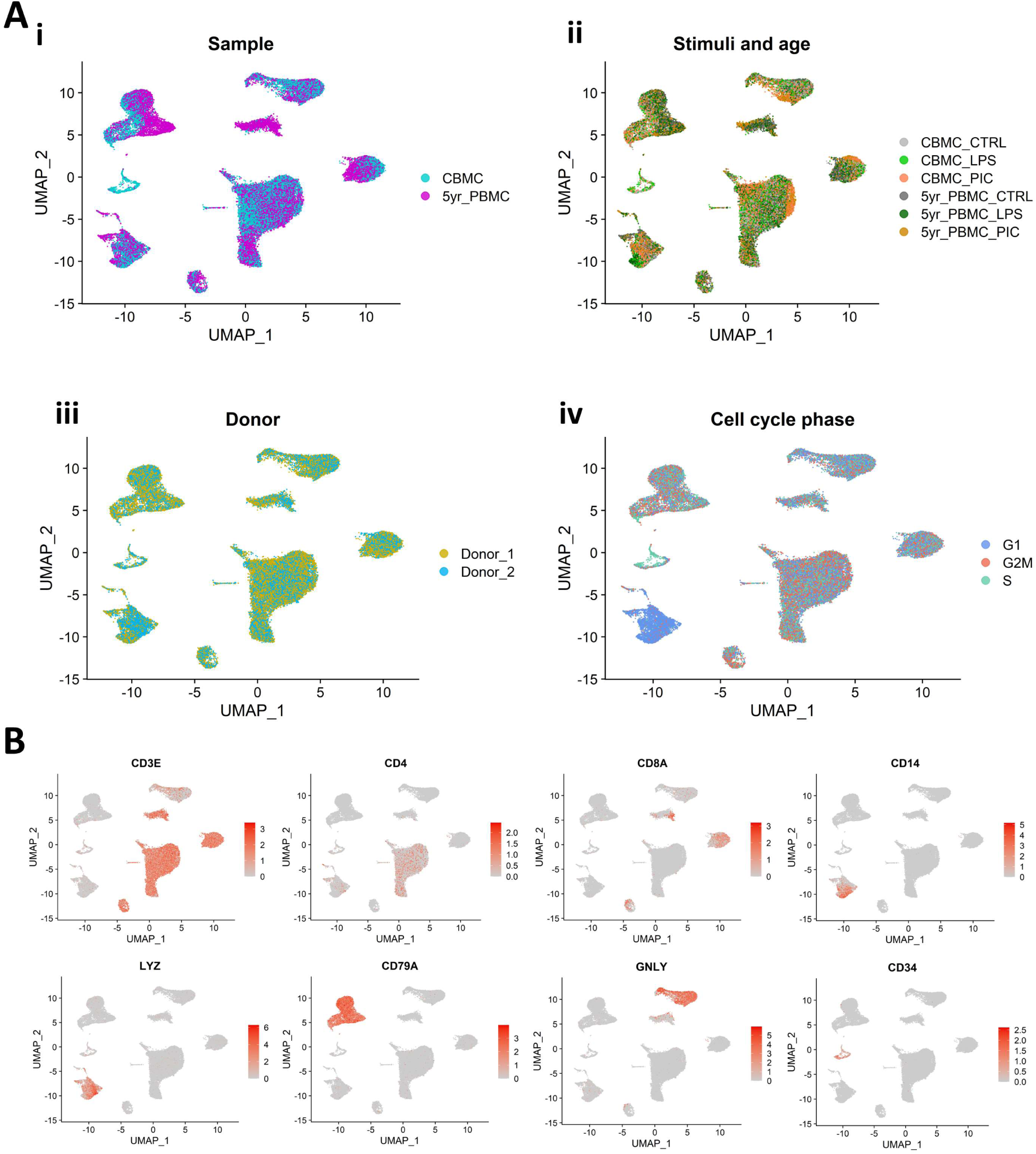
Integrated UMAP plots overlayed with selected sample characteristics. **A**) UMAP plots stratified by sample collection time point (**i**), stimuli/age group (**ii**), Biological donor (**iii**), and cell cycle phase (**iv**). **B**) Integrated UMAPs overlayed with selected marker genes.

**Figure S2.**
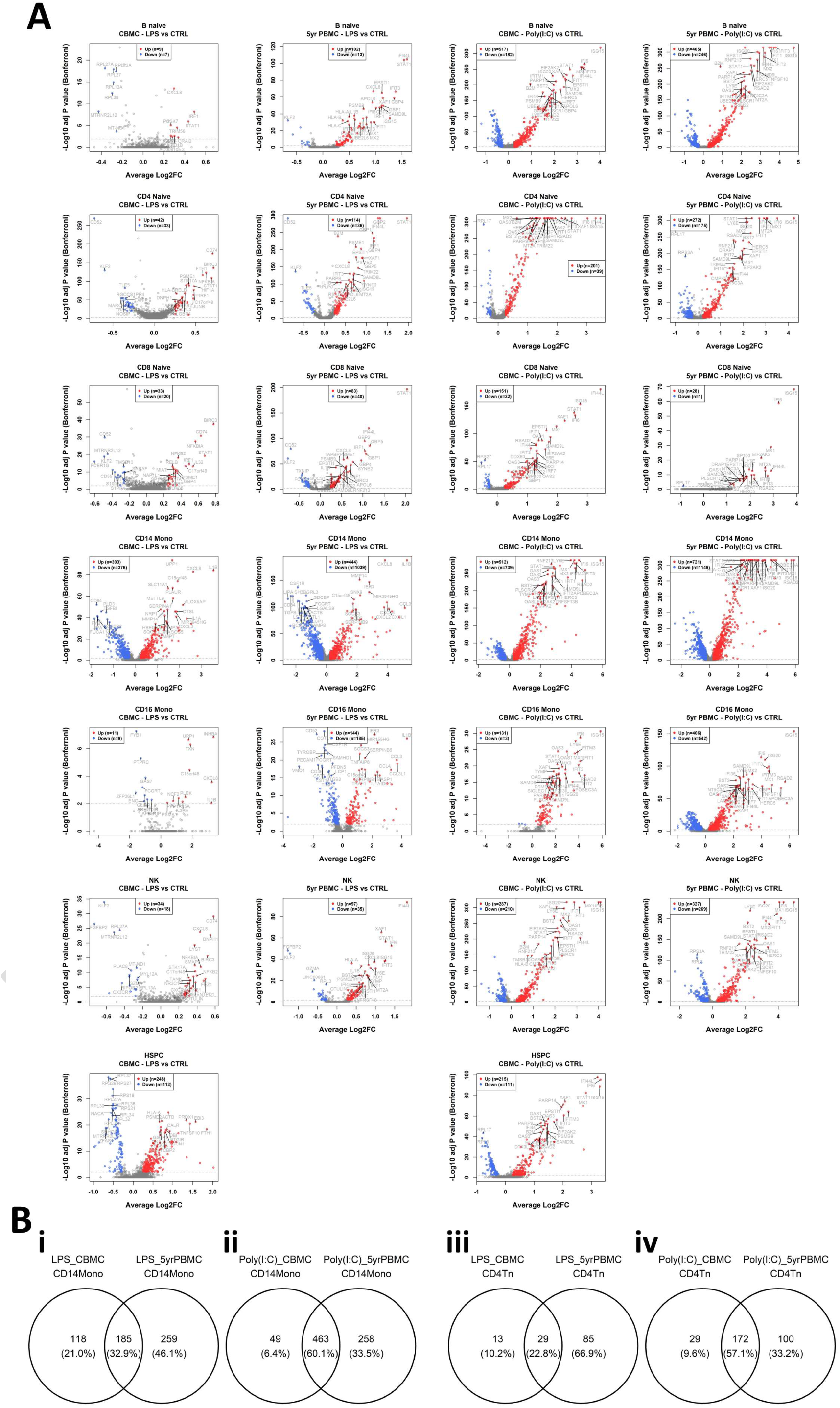
**A**) Volcano plots of selected comparisons from the differential expression analysis. Each point represents a gene, and the x-axis shows the average Log_2_-fold change, and the y-axis shows the -Log_10_ Bonferroni-corrected *p* value for the corresponding comparison. Genes which recorded a corrected *p* value < 0.01 and an absolute average log_2_-FC > 0.25 were considered significantly dysregulated and are shown as red (upregulated) and blue (downregulated) points. The total number of up- and down-regulated genes are shown in the inset of each plot, with selected dysregulated genes annotated. **B**) Venn diagrams showing the overlap of significantly upregulated genes between CBMC and 5yr PBMC samples for LPS (**i**) and Poly(I:C) (**ii**) stimulated CD14^+^ Monocytes and LPS (**iii**) and Poly(I:C) (**iv**) stimulated naïve CD4^+^ T cells.

**Figure S3.**
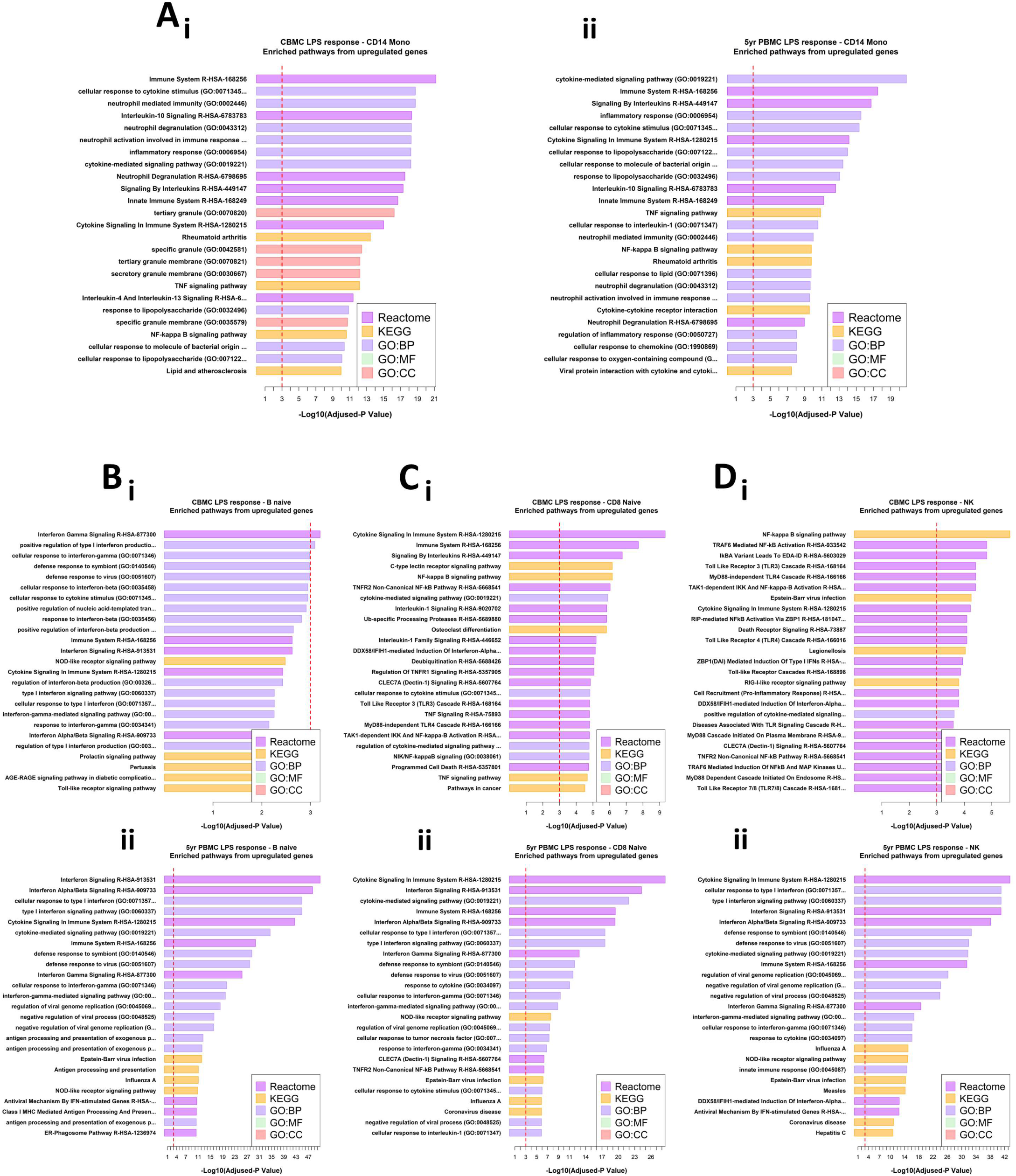
Pathways analysis of upregulated genes identified by differential expression analysis for selected cell types stimulated with LPS versus their corresponding unstimulated sample. Horizontal bar plots of the top 25 significantly enriched pathways found for the comparison of CD14^+^ monocytes (**A**), Naïve B cells (**B**), Naïve CD8^+^ T cells (**C**), and NK cells (**D**) for CBMC (**i**) and 5yr PBMC (**ii**) samples. The panel in A runs left to right and the panes in B-D run top to bottom. The x-axis shows the -Log_10_ adjusted-P value associated with pathways enrichment, the dashed red line indicates an adjusted-*p* value of 0.001. Results are ordered from top by decreasing adjusted-*p* value for significantly enriched pathways identified from the Reactome, KEGG, and Gene Ontology (GO) databases. BP, Biological Process; MF, Molecular Function; CC, Cellular Compartment. See **supplementary data** for complete lists of significantly enriched pathways from dysregulated genes for all cell types analysed.

**Figure S4.**
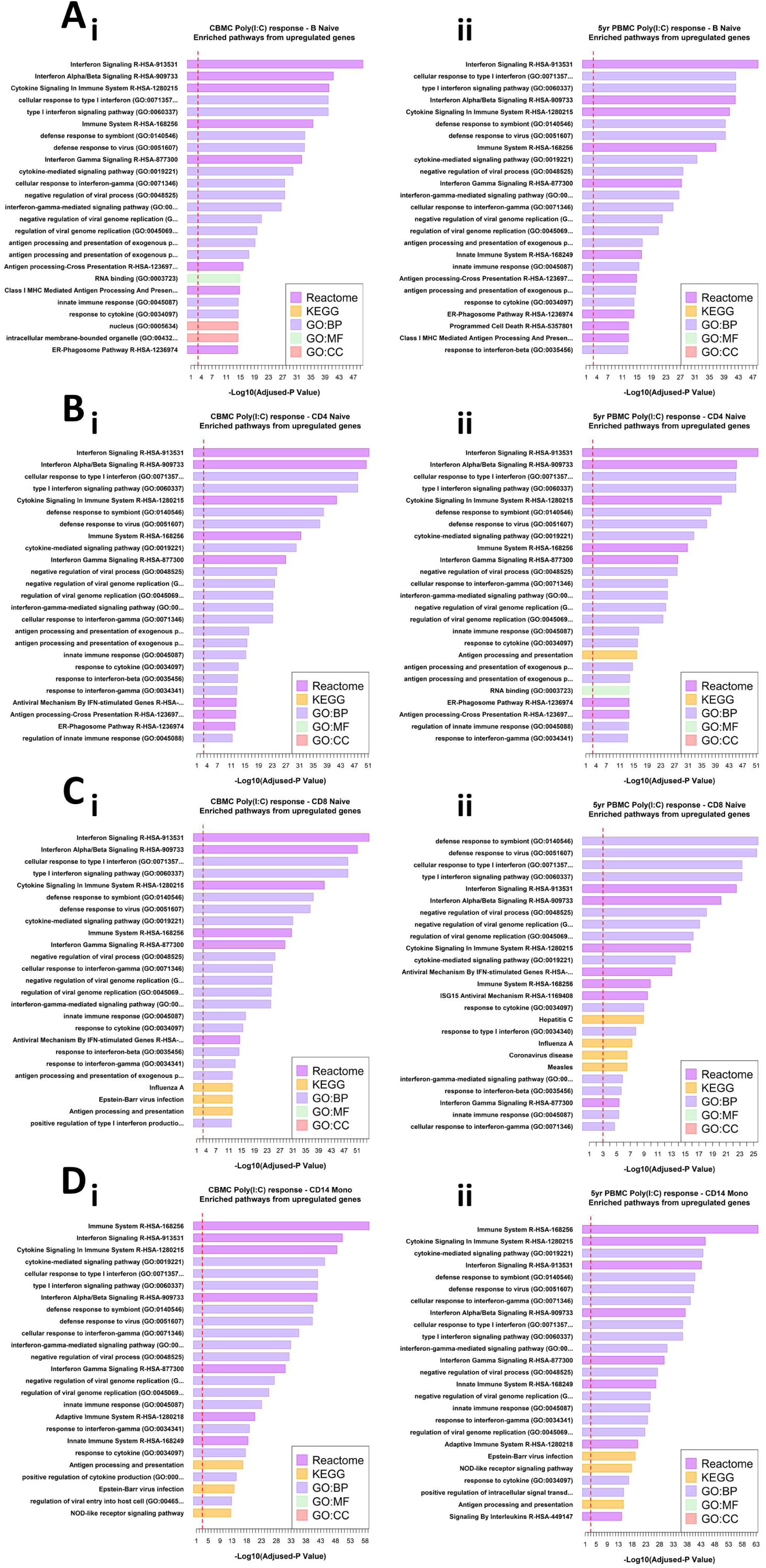
Pathways analysis of upregulated genes identified by differential expression analysis for selected cell types stimulated with Poly(I:C) versus their corresponding unstimulated sample. Horizontal bar plots of the top 25 significantly enriched pathways found for the comparison of Naïve B cells (**A**), Naïve CD4^+^ T cells (**B**), Naïve CD8^+^ T cells (**C**), and CD14^+^ Monocytes (**D**) for CBMC (**i**) and 5yr PBMC (**ii**) samples. The x-axis shows the -Log_10_ adjusted-*p* value associated with pathways enrichment, the dashed red line indicates an adjusted-*p* value of 0.001. Results are ordered from top by decreasing adjusted-*p* value for significantly enriched pathways identified from the Reactome, KEGG, and Gene Ontology (GO) databases. BP, Biological Process; MF, Molecular Function; CC, Cellular Compartment. See **supplementary data** for complete lists of significantly enriched pathways from dysregulated genes for all cell types analysed.

**Figure S5.**
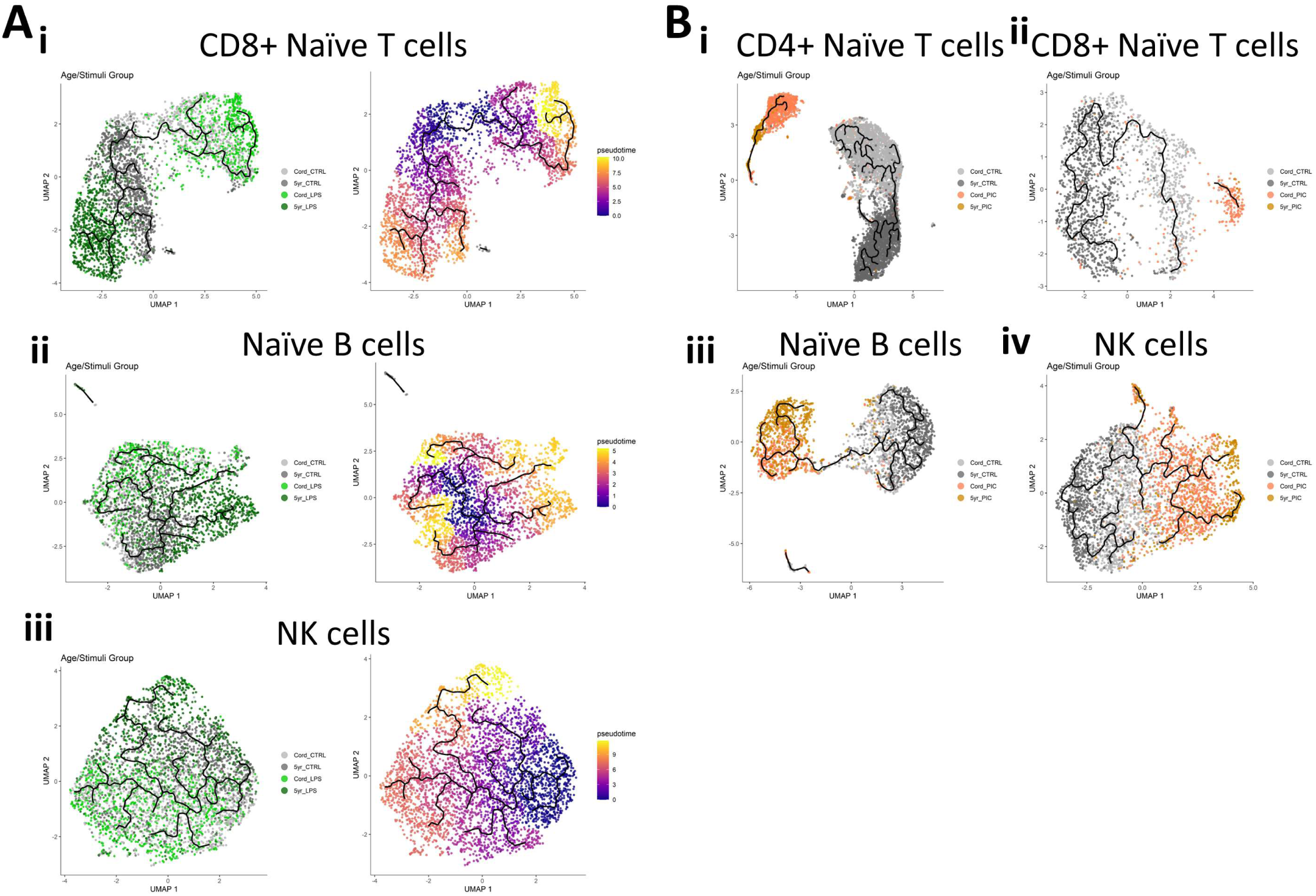
Monocle3 pseudotime trajectory analysis results for selected cell types. UMAP plots representing cell activation trajectories for LPS-stimulated (**A**) CD8^+^ naïve T cells (**i**), naïve B cells (**ii**), and NK cells (**iii**), and Poly(I:C)-stimulated (**B**) naïve CD4^+^ T cells (**i**), naïve CD8^+^ T cells (**ii**), naïve B cells (**iii**), and NK cells (**iv**). In panel **A**, the first plot (**i**) in each panel is stratified by sample group and the second plot (**ii**) displays pseudotime. The branching black line on each plot represents the activation trajectory fitted to the data.

**Figure S6.**
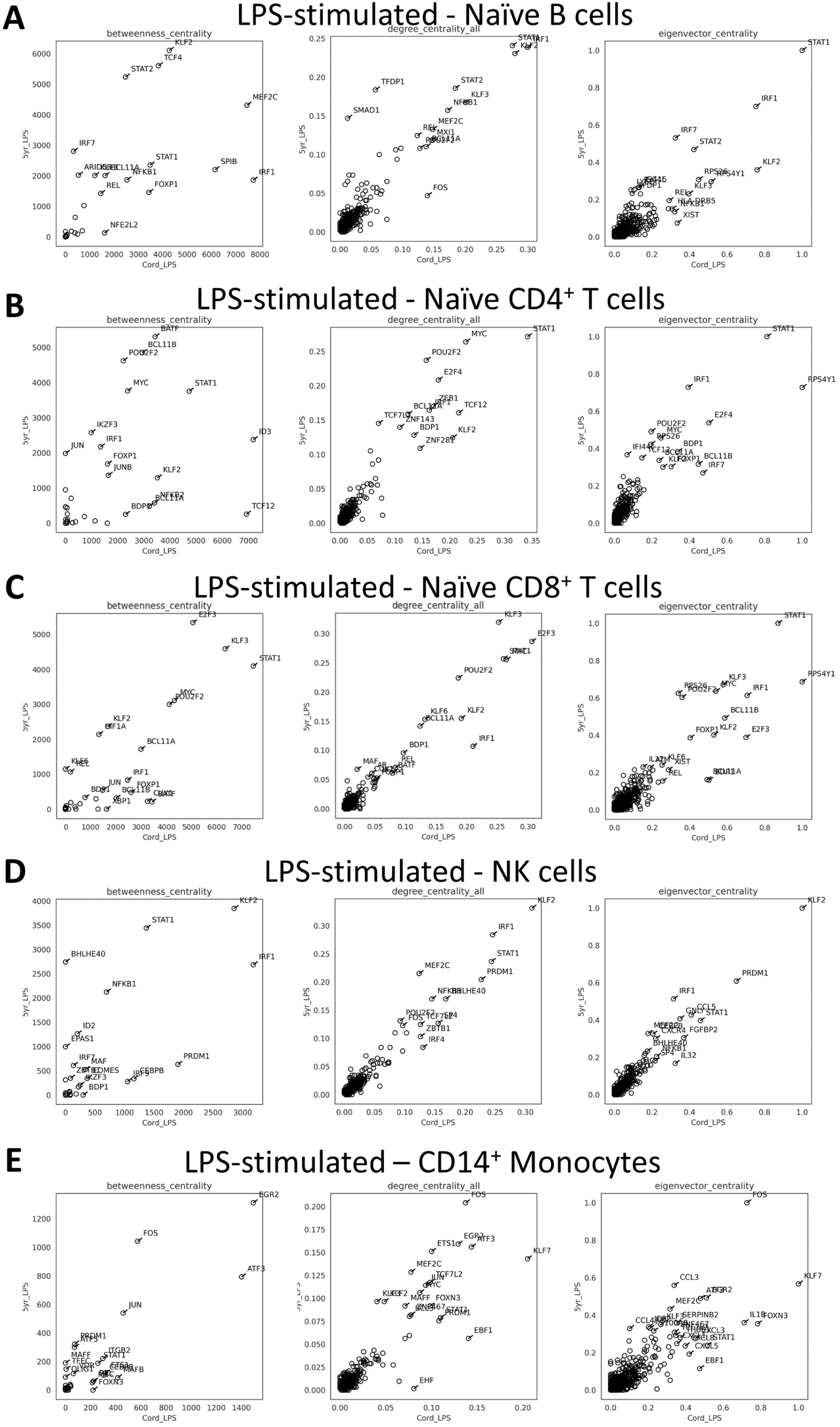
Gene Regulatory Network characteristics for selected LPS-induced cell types. Plots show the correlation between the metrics for transcription factors and target genes for samples collected from cord blood (CBMC) (x-axis) and 5yr blood (PBMC) (y-axis). Plot are included for naïve B cells (**A**), naïve CD4^+^ T cells (**B**), naïve CD8^+^ T cells (**C**), NK cells (**D**), and CD14^+^ monocytes (**E**), and show the betweenness centrality (left), degree centrality (middle), and eigenvector centrality (right). In each case, a higher value indicates a greater importance within the network and a deviation off the diagonal indicates the metric has a higher value in the corresponding sample.

**Figure S7.**
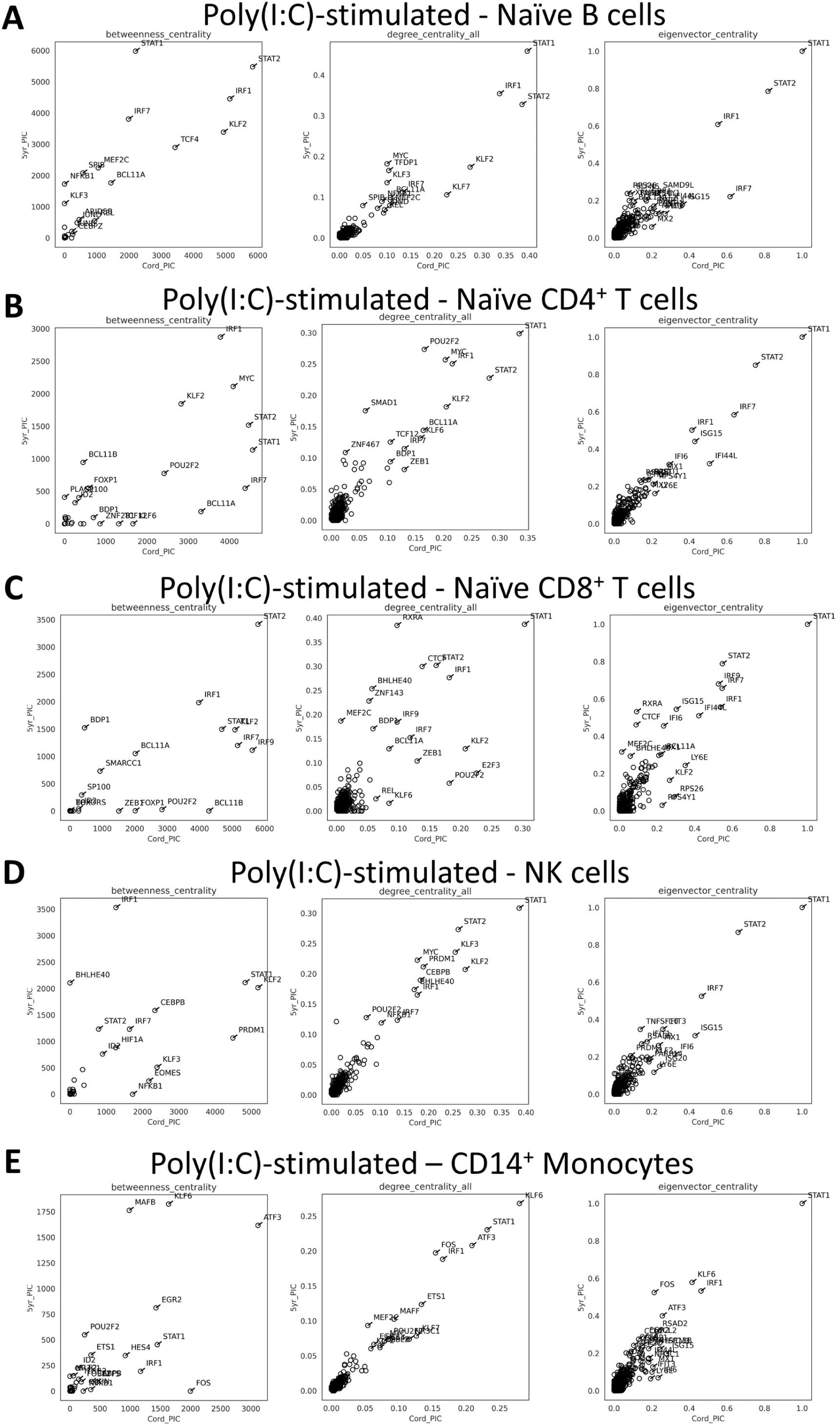
Gene Regulatory Network characteristics for selected Poly(I:C)-induced cell types. Plots show the correlation between the metrics for transcription factors and target genes for samples collected from cord blood (CBMC) (x-axis) and 5yr blood (PBMC) (y-axis). Plots are included for naïve B cells (**A**), naïve CD4^+^ T cells (**B**), naïve CD8^+^ T cells (**C**), NK cells (**D**), and CD14^+^ monocytes (**E**), and show the betweenness centrality (left), degree centrality (middle), and eigenvector centrality (right). Plot and metric characteristics are the same as above (Figure S6).

**Figure S8.**
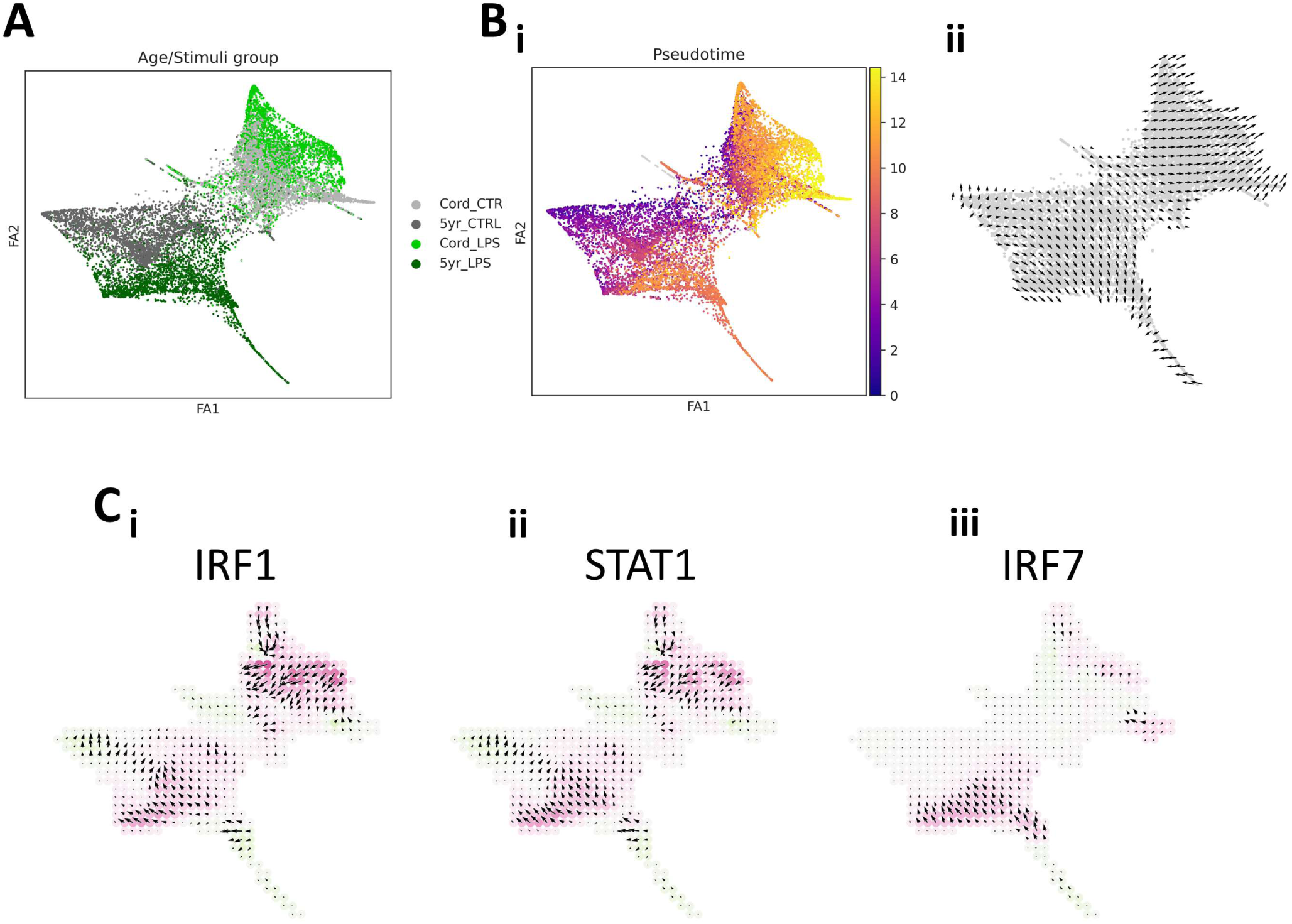
*In silico* perturbation to nullify transcription factor activity (CellOracle). **A**) Force-directed graph of naïve CD4^+^ T cells stratified by stimuli/age group. This plot recapitulates the characteristics observed from the UMAP (Figure 3B(i)). **B**) **i**) Monocle-defined pseudotime showing differentiation into CBMC-related and 5yr PBMC-related activation states and **ii**) differentiation/activation vectors projected onto the force-directed graph (PAGA). **C**) Knock-out (KO) simulation vector field with perturbation scores for *in silico* KO of *IRF1* (left), *STAT1* (center), and *IRF7* (right). KO of *STAT1* and *IRF1* results in remarkably similar effect on cellular identity. Colors correspond to the perturbation score and denote whether the KO would putatively block (red) or promote (green) activation on that region of the vector field. Arrows indicate the directional change in activation following simulated KO and are calculated as the inner product of the activation trajectory (pseudotime) and the simulated perturbation score. The size of the arrow represents the magnitude of the inner product.

## Extended Methods

### Materials and Methods

#### Study subjects

The study was designed to assess matched birth (CBMC) and 5 years (PBMC) blood samples following LPS and Poly(I:C) treatment, along with matched untreated controls, from two donors (one male, one female). The samples were curated from the Childhood Asthma Study (CAS) cohort, a prospective birth cohort for high risk of asthma development(1–5). Cord blood samples were collected from healthy, full-term, singleton births. Matched 5-year samples were collected from the same donor by home visit close to their 5th birthday. This study has ethics approval by The University of Western Australia (reference RA/4/1/7560), and fully informed parental consent was obtained for sample collected from each subject.

#### In vitro cell culture and innate immune stimulation

Cryopreserved CBMC/5yr PBMC samples were thawed in RPMI 1640 (Gibco) + 10μl DNase (1st freeze/thaw cycle), washed (centrifuged at 400g for 7 mins at RT) and resuspended in 1ml PBS + 2% AB serum (heat inactivated). Residual red blood cells were depleted from cord blood samples with an EasySep RBC Depletion kit (Stemcell Technologies), as per the manufacturer’s protocol. Cell viabilities are recorded in Table S1. Cells were resuspended at 1×106 cells per ml in RPMI + 5% AB serum and 0.25×106 cells were transferred to dedicated wells of round bottom 96-well polystyrene culture plates (Thermo Fisher Scientific). Four wells (∼1×106 cells) were allocated to each condition. Wells were stimulated with 1ng/ml LPS (Enzo Biochem, Cat No. ALX-581-007-L001, derived from E. coli, serotype R515) or 50μg/ml Poly(I:C) (InvivoGen, Cat. Code: tlrl-pic) or left untreated, and plates were incubated at 37oC (5% CO2) for 18 hours. LPS is a bacterial cell wall component and the quintessential TLR4 ligand. Poly(I:C) is a synthetic analogue of double-stranded RNA (dsRNA) and a potent activator of TLR3 and other nucleic acid sensing receptors (e.g., RIG-I, MDA-5)(6). Each cryopreserved vial was processed on a different day, so that matched stimuli/control samples were processed together. Following culture, replicate culture plate wells (4 per sample/condition) were gently resuspended and transferred to a sterile 1.5ml LoBind tube (Eppendorf). Samples were pelleted (centrifugation at 500 x g for 7 minutes at 4oC) and re-suspended in PBS + 0.04% BSA (UltraPure; Thermo Fisher Scientific) (4oC) to a target concentration of 2,000 cells/μl. Post-culture viability is recorded in Table S1. Samples were immediately transferred on ice to Genomics WA (Perth, Western Australia) for library preparation and sequencing.

#### Alignment and initial quality control

Raw fastq.qz files were processed with the CellRanger Toolkit (Version 6.1.1, 10x Genomics) with the Human GRCh38 genome assembly (refdata-gex-GRCh38-2020-A) was used as the reference genome. The CellRanger count pipeline was run with default parameters. CellRanger web_summary outputs were assessed and no alerts (warnings or errors) were recorded for any sample. Selected CellRanger outputs are recorded in Table S1; briefly, this project generated (on average) 5,527.17 cells per sample with 63,098.75 mean reads per cell and an average of 1980.17 genes detected per cell, as estimated by CellRanger. The raw feature matrix, and corresponding barcodes and features, were used for downstream QC and analysis.

#### Sample pre-processing and quality control

Count matrices and corresponding barcode and feature files were imported into the R statistical environment (version 3.6.2), and all subsequent QC/analyses were conducted in R, unless otherwise stated. Each sample was run through a QC pipeline which combines functions from several R packages, of which Version 3.2.0 of Seurat(7) was used most extensively for QC. Cells were excluded if they had a mitochondrial gene content greater than three median absolute deviations (MADs) above the median or a ribosomal gene content > 50% (which excluded few cells). Features were filtered to only those that were expressed in at least 0.01% of cells. Cells with low feature counts (approximately < 2000) were also excluded, according to dynamic thresholding of the count distribution. Doublets were detected and removed with DoubletFinder(8) and the cell cycle phase was estimated with the CellCycleScoring function using known cell cycle related genes (Seurat). Pre-processing and quality control metrics are recorded in Table S1. Raw and processes datasets are available via the Gene Expression Omnibus (GSE232186).

#### Integration, Annotation, and Dimensionality reduction

Individual pre-processed samples were integrated with Seurat(7) using default parameters. Individual cells were annotated with Azimuth(9) using the human PBMC reference data set. Level 2 cell type resolution (default) was used, and cell were excluded if they had a score < 0.5. Dimensionality reduction (UMAP) plot coordinates from the Azimuth reference were used to show the region/cell type that corresponds to our scRNA-Seq data, and a UMAP plots were also generated from the integrated data with Seurat (e.g., ScaleData, RunPCA, and RunUMAP functions) to display the integrated cell type clustering, as well as group variables and marker gene expression intensities.

#### Differential gene expression and Pathways analysis

To identify differentially expressed genes (DEGs) between LPS/Poly(I:C) and corresponding unstimulated control samples for each cell type, we employed MAST(10) via the FindMarkers function from Seurat, and included cellular detection rate, mitochondrial gene proportion, and cell cycle phase. As each donor represents a different biological sex (male, female), this variable was also included as a latent variable. This approach was selected to accommodate our small sample size (two biological donors), although we acknowledge that incorporating individual participant variation as a latent variable is a suboptimal approach compared to other methods suited to larger sample sizes (viz. pseudobulk and mixed models with a random effect for individual)(11,12), and this is a limitation of our study. However, our analysis identified many co-regulated genes which would be expected to be dysregulated following treatment with LPS and Poly(I:C) (corroborated by pathways enrichment analysis and independently identified with GRN analysis) and the primary genes of interest recorded extremely small Bonferroni-corrected p values (generally < 10×10-50). For these reasons, we believe the loss in precision of the MAST + latent variable method had limited impact on the findings in our study. Genes were considered differentially expressed if they recorded a Bonferroni-corrected p value < 0.01 and an average Log2 fold-change in expression of > 0.25 (upregulated) or < -0.25 (downregulated). Aligned volcano plots displaying DEGs identified from multiple cell type simultaneously were plotted in base R and DEG count heatmaps were plotted with the pheatmap function in R. Significantly enriched pathways associated with DEGs between cell type/stimuli groups were identified with enrichR(13) in R to query biologically-relevant annotated gene sets from the Reactome(14), KEGG(15), and Gene Ontology(16) databases.

#### Pseudotime trajectory inference

We applied Monocle3(17) to infer stimuli-related activation trajectories from transitional cellular states present in the data. For each analysis, only the raw counts from cells relevant for that comparison (e.g., CBMC/5yr PBMC untreated and LPS-treated monocytes) were included. Default parameters were used for pre-processing and the data was aligned with the donor variable as alignment group and the cellular detection rate and mitochondrial content included as terms in the model formula. UMAP dimensionality reductions were run with 10 nearest neighbours and a minimum distance of 0.1 and cells were clustered with a k value (nearest neighbours) of 10 and a resolution value of 0.001 for all comparisons, with the exception of cluster resolution values of 0.002 and 0.005 for LPS- and Poly(I:C)-induced naïve CD4+ T cells comparisons, respectively, to accommodate larger cell numbers. Regions enriched with unstimulated controls were selected as pseudotime start points so that trajectories extended into stimuli-activated regions.

#### Gene Regulatory Network (GRN) analysis and in silico perturbations

We employed CellOracle(18) to build GRNs in order to identify the key molecular drivers (transcription factors (TF)) and their corresponding target genes for selected cell type/stimuli groupings. For this analysis, SCANPY(19) (version 1.9.3) was used for pre-processing and force directed graph construction (Partition-based graph abstraction (PAGA)(20)) and CellOracle (version 0.12.0) was run with Python (3.10.6) on Ubuntu 22.04.1 via Windows Subsystem for Linux 2 kernel. Genes with at least 1 count were retained and then genes were filters to identify the top 3000 most variable for each comparison. The data was normalized (scanpy.pp.normalize_per_cell), log transformed (scanpy.pp.log1p) and scaled (scanpy.pp.scale) with default parameters. Additionally, the donor variable was adjusted for in the data with scanpy.pp.regress_out. Separate analyses were run from raw counts for each cell type/stimuli comparison and group specific GRNs (e.g., Cord_Control, Cord_LPS, 5yr_Control, and 5yr_LPS) were constructed from the Human promoter base GRN provided. CellOracle was run with standard parameters and the monocle3-defined pseudotime values for each cell were included for analysis. From the output of CellOracle, we plotted Venn diagrams (ggvenn R package) to show the overlap of significantly connected (p value < 0.01) target genes (TG) of IRF1, IRF7, STAT1, and STAT2 for each comparison, and displayed TF-TG wiring diagrams of the top 100 TG connections with the igraph R package. CellOracle constructed GRNs were then used to perform in silico transcription factor perturbations of IRF1, IRF7, and STAT1 to simulate the changes in cellular states after nullifying the regulatory effects of the TFs. The scale parameter was adjusted to suit each comparison (as recommended) and standard parameters were used for all other functions in this analysis.

#### Ligand-Receptor interaction analysis

We employed CellCall(21) to identify putative ligand-receptor (L-R) communication between selected cell types following LPS- and Poly(I:C)-induced activation. For each stimuli/age comparison, the genes were filtered to the top 3000 most variable (compared to corresponding unstimulated control samples) and single cell profiles were restricted to selected cell types of interest. Raw counts were use as input, the transcriptional communication profile was calculated with default parameters, and L-R interactions stratified by cell type were visualized with circus plots. The LR2TF function was used to extend the analysis by capturing putatively activated TFs downstream of receiver cell receptor binding of sender cell ligands for communication between CD14+ monocytes and naïve CD4+ T cells.

